# Understanding the regulation of chronic wounds by tissue inhibitors of matrix metalloproteinases through mathematical modelling

**DOI:** 10.1101/2024.09.20.614051

**Authors:** Sonia Dari, Reuben D. O’Dea, Nabil T. Fadai

## Abstract

Understanding the biochemistry and pharmacodynamics of chronic wounds is of key importance, due to the millions of people in the UK affected and the significant cost to the NHS. Chronic wounds are characterised by elevated concentrations of matrix metalloproteinases (MMPs) that destroy the surrounding extracellular matrix (ECM). However, fibroblasts can produce tissue inhibitors of MMPs (TIMPs) in order to regulate wound healing. Therefore, the role of TIMPs in both acute and chronic wounds needs to be properly understood in order to develop therapeutic treatments. In this work, we propose a reaction-diffusion system of four partial differential equations that describe the interaction of the ECM, fibroblasts, MMPs, and TIMPs in a wound. We observe that, subject to biologically realistic parameter values, this mathematical model gives rise to travelling wave solutions. Using bifurcation analysis, we demonstrate that deregulated apoptosis in the ECM results in the emergence of chronic wounds, and the reversal of these chronic wounds is prohibited for lower TIMP production values. These results are replicated within a simplified model obtained via a parameter sensitivity analysis. This model is further extended to more realistic spatial domains where we demonstrate the effectiveness of a therapeutic hydrogel containing TIMPs as a treatment for chronic wounds.

## 1 Introduction

Wound healing is a physiological process that involves the coordinated action of various cell types and biochemical agents to repair tissue damage [1, 2]. In the UK, approximately 2.2 million people suffer from so-called “chronic wounds”. These wounds usually last 12–13 months and recur in 60%–70% of cases, potentially leading to loss of function, and costing the NHS over £5 billion annually [3, 4]. This issue is particularly significant in diabetes patients, who are more prone to developing chronic wounds that require medical intervention to aid healing [5, 6]. Understanding the biochemical mechanisms underpinning chronic wounds is essential for developing new treatments.

A wound is defined as damage to the two layers of the skin: the epidermis and the dermis. The epidermis, the outermost layer, serves as a barrier against infection, while the dermis, the innermost layer, gives the skin tensile strength through the dermal extracellular matrix (ECM) [7, 8]. Damage to the skin tissue results in a wound, initiating the four stages of the wound healing process: haemostasis, inflammation, proliferation and remodelling [1]. A wound that successfully progresses through these four stages in a well-coordinated manner is defined as an acute wound, whereas a wound that persists for a prolonged time in any of these four stages is classified as a chronic wound [5].

During the inflammation and proliferation stages of wound healing, various cell types including keratinocytes, fibroblasts, endothelial cells, and inflammatory cells produce matrix metalloproteinases (MMPs) [5, 9]. MMPs facilitate the breakdown of protein components within the extra-cellular matrix (ECM), enabling fibroblast migration and promoting cell proliferation and tissue modification [5]. Moreover, an increased expression of MMPs is observed at the edge of the healing wound throughout all stages of acute wound healing [10]. Maintaining a properly regulated spatio-temporal concentration of MMPs during this process is crucial. Low MMP levels can lead to uncontrolled ECM production, potentially resulting in issues such as hypertrophic scarring or dermal fibrosis [11]. Conversely, excessive MMP levels can lead to chronic wounds, as the heightened MMP activity leads to increased ECM degradation [12]. One reason for these increased MMP levels is the deregulated apoptosis of dermal tissue, which can be triggered by uncontrolled blood sugar levels in patients with diabetes [13, 14]. In response to these elevated MMP concentrations, tissue inhibitors of MMPs (TIMPs) are produced, primarily by fibroblasts. TIMPs regulate the activity of MMPs by binding to them, thus inhibiting their ability to degrade the ECM and in doing so, act as regulators of ECM remodelling and cellular behaviour [15].

Modulating the activity of TIMPs has been shown to influence wound healing outcomes, suggesting their potential as therapeutic agents [16–18]. One approach to deliver TIMPs effectively is through hydrogel therapies. Hydrogels are three-dimensional hydrophilic polymer networks with various applications in the soft tissue engineering of blood vessels, muscles and skin. They are composed of either physical or chemical cross-linked bonds, designed to mimic the microenvironment of skin via their porous and hydrated structure [19]. Hydrogels can also contain bioactive molecules which, once placed on the a tissue surface, deliver these bioactive substances with spatial and temporal control [19]. Therefore, utilising hydrogels to deliver TIMPs could provide a targeted and controlled method for enhancing wound healing, potentially offering an effective treatment for chronic wounds.

Employing mathematical models to explore the wound healing process offers a valuable framework for understanding chronic wounds and may potentially lead to advancements in the development of new treatments, thus enhancing patients’ quality of life. Continuous mathematical models representing both dermal and epidermal wound healing commonly adopt the structure of reaction-diffusion systems. These models build upon the research of Sherratt and Murray [20], who investigated the interplay between epidermal cells and a representative chemical serving as a regulator of mitosis. Their model has subsequently been extended to consider the role of other chemical agents involved in epidermal wound healing [21–24] as well as the restoration of the ECM in dermal wound healing, such as through growth factors [25, 26] and the role of angiogenesis [7, 27– 33]. Other models adopt a hybrid multiscale approach to consider the simultaneous healing of the epidermis and the dermis [34–36] as well as to consider the role of collagen fibre orientation due to cell movement in wound healing [37–41]. More recent advances include mechanochemical modelling to consider the role of ECM contraction in wound healing [35, 42–46]. We note that many of the aforementioned mathematical models display travelling wave solutions, aligning with experimental findings from wound healing assays [47, 48]. While the models discussed above primarily address the healing of acute wounds, some research has shifted focus to the prolonged inflammatory phase characteristic of chronic wounds [49–54], as well as effects of various treatments on such chronic wounds [54–57].

By integrating both acute and chronic wound models, a more comprehensive understanding of wound healing dynamics can be achieved. Our previous work [18] has considered the interaction of MMPs and dermal cells in the presence of both acute and chronic wounds, giving rise to travelling wave solutions in both parameter regimes. Through direct simulations, travelling wave analysis and bifurcation analysis, we demonstrate that deregulated apoptosis results in the emergence of chronic wounds, characterised by elevated concentrations of MMPs. As well this, we observe a hysteresis effect when the apoptotic rate is varied, indicating multistable travelling waves depending on whether we initially begin from an acute or a chronic wound, giving insights into the management and potential reversal of chronic wounds.

Although extensive research into wound healing has been conducted, the literature currently lacks a thorough exploration of TIMPs as regulators in both acute and chronic wounds, and their potential as therapeutic treatments for chronic wounds. In this work, we extend the model framework presented in [18] by examining the role of TIMPs as regulators of MMP concentrations and hence as regulators of chronic wounds. In Section 2, we develop our mathematical model which we then nondimensionalise in Section 2.1. We present numerical simulations of the model in Section 2.2 which demonstrates the existence of travelling wave solutions, thus motivating a steady state and bifurcation analysis in Section 3. In Section 4, we conduct a parameter sensitivity analysis and exploit this to obtain a reduced model that we then demonstrate supports the same qualitative dynamics as the original model. In Section 5, we extend the reduced model to two dimensions. Here, we demonstrate that elevated MMP production levels lead to the emergence of a chronic wound. Furthermore, we demonstrate how treating such a chronic wound with a TIMP-containing hydrogel applied to the wound surface, modelled using a Robin boundary condition, can reverse chronic wounds. Finally, we discuss our results in Section 6.

## 2 Model Development

We present a mathematical formulation to describe the dynamics of MMPs and TIMPs within a wound and consequently, the impact on the restoration of the ECM by fibroblasts. Elevated concentrations of MMPs play a key role in the emergence of chronic wounds, while TIMPs are produced in order to regulate MMP concentrations. We construct a reaction-diffusion model to capture these dynamics in the wound healing process. For simplicity, we consider a wound in a one-dimensional Cartesian geometry, where *x* denotes the longitudinal distance across a fixed domain [0, *L*], and we examine the wound’s evolution within this domain over time *t*. This model consists of four populations: ECM density, denoted as *c*(*x, t*); fibroblast cell population density, *f* (*x, t*); MMP concentration, *m*(*x, t*) and TIMP concentration, *p*(*x, t*). Our model seeks to include all the key processes involved in the interactions between these populations in the context of a wound (see Figure 1), resulting in a complex system with numerous parameters. In Section 4, we will use a parameter sensitivity analysis to reduce the complexity of the model.

**Figure 1:**
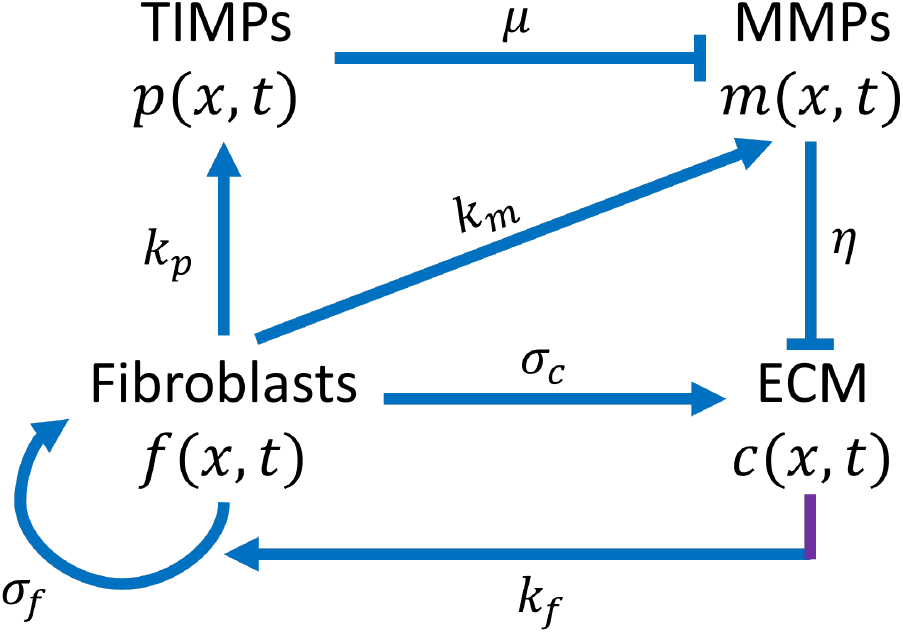
Schematic of the interactions of ECM, fibroblasts, MMPs and TIMPs

We assume that the components of the ECM are produced by fibroblasts with production rate *σ*_*c*_, which decreases to zero as the ECM concentration approaches the carrying capacity *c*_0_, noting that *c* = 0 corresponds to the complete absence of ECM. As mentioned in Section 1, MMPs can assist in the healing of the wound up to a threshold concentration, denoted here as *m*_thresh_, above which they have zero healing contribution. This effect is incorporated in the production process. The degradation of the ECM by MMPs occurs with rate *η*. Furthermore, the ECM undergoes apoptosis with rate *δ*_*c*_ and its motion within the wound is represented by linear diffusion with effective diffusivity *D*_*c*_.

Fibroblasts undergo mitosis: they are produced by local connective tissue and reproduce by logistic growth. Moreover, fibroblasts undergo apoptosis with rate *δ*_*f*_, linear diffusion with effective diffusivity *D*_*f*_, and migrate chemotactically towards the absence of healthy tissue, with rate *χ*_*f*_, in response to various chemical signals such as growth factors and cytokines at the wound site, which arise from various cell types such as fibroblasts and keratinocytes.

MMPs are produced by fibroblasts in response to inflammatory chemicals which are found in large quantities in a wound. MMPs diffuse through the wound with diffusivity *D*_*m*_ and decay with rate *δ*_*m*_. MMPs are inhibited by TIMPs and thus become inactive, leading to the effective removal of MMPs with rate *µ*. TIMPs are produced by fibroblasts in response to the presence of MMPs, decay with rate *δ*_*p*_, and are also depleted in the process of MMP inhibition with rate *µ*.

Combining these aforementioned assumptions yields the following set of PDEs describing the spatiotemporal evolution of *c*(*x, t*), *f* (*x, t*), *m*(*x, t*) and *p*(*x, t*):

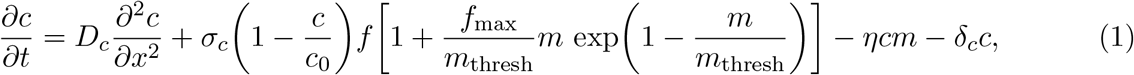

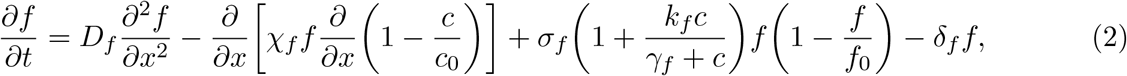

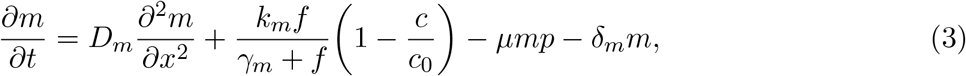

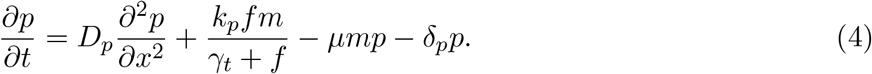

The initial conditions are chosen to model a symmetric wound beginning at 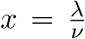(with *ν* being a dimensionless constant). Specifically, within healthy tissue 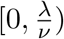, ECM and fibroblasts are initially at carrying capacity (*c*_0_ and *f*_0_ respectively) and absent within the wound region 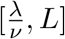. We also assume that a small spike in MMPs is initially present at the wound edge while TIMPs are absent throughout. These modelling assumptions yield the following initial conditions:

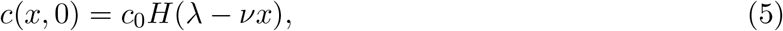

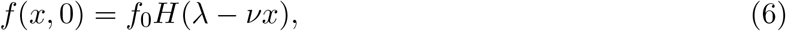

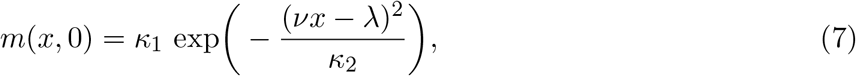

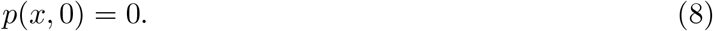

In (5)–(6), *H* is the Heaviside function, while *c*_0_ and *f*_0_ correspond to the carrying capacity of the ECM and fibroblasts, respectively. Moreover *κ*_1_ represents the peak value of MMPs and *κ*_2_ corresponds to the spike width of MMPs in (7). Since we assume the wound to be symmetric and the surrounding tissue to be healed far away from the wound, we adopt zero-flux boundary conditions on both ends of the finite spatial domain [0, *L*]:

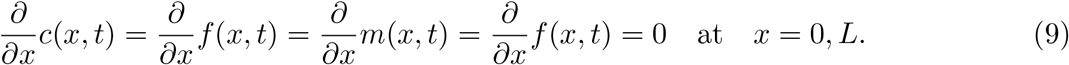

### 2.1 Non-dimensionalisation

We non-dimensionalise the system (1)–(9) in order to reduce the number of parameters and thus simplify the analysis of the system. Since we are interested in the behaviour of the system on the timescale of production rate of the ECM, we introduce the following dimensionless variables:

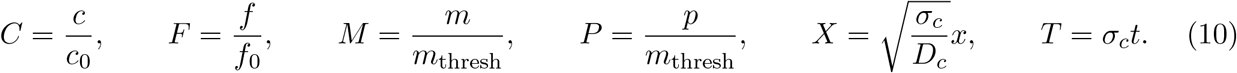

Applying these scalings to (1)–(4) yields the following dimensionless equations:

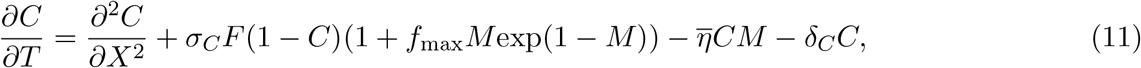

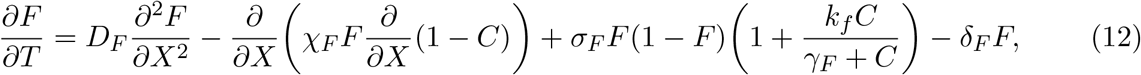

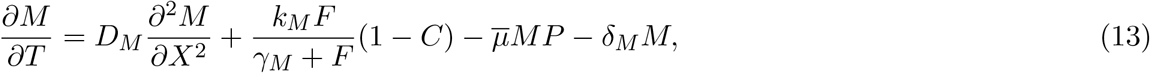

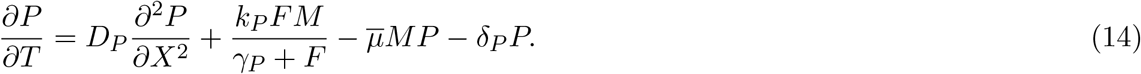

For simplicity, we take 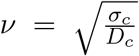 in (5)–(7) and *κ*_2_ = 1 in (7) which leads to the following nondimensional initial and boundary conditions:

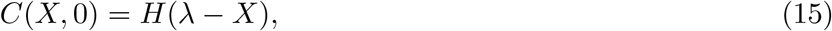

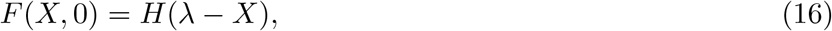

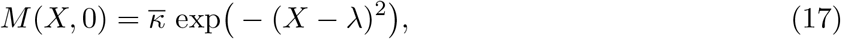

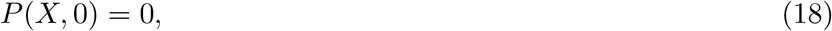

and

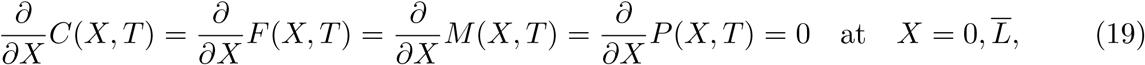

where 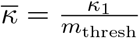 and 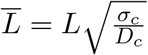. We remark that the dimensionless values 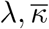 and 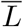 are chosen for demonstrative purposes and the corresponding dimensional parameter values are straightforward to calculate and align with physiological values.

We denote the dimensionless system (11)–(19) as the ‘regulated wound healing model’ (RWHM). Suitable values for the dimensional parameters in (1)–(4) are given in Table 1 and represent healthy biological functioning, noting that the parameters that were unable to be obtained from biological literature were estimated heuristically. The dimensionless parameter values in the RWHM are given in Table 2. We note that unless otherwise stated, these are the parameter values used in all subsequent numerical simulations.

**Table 1:**
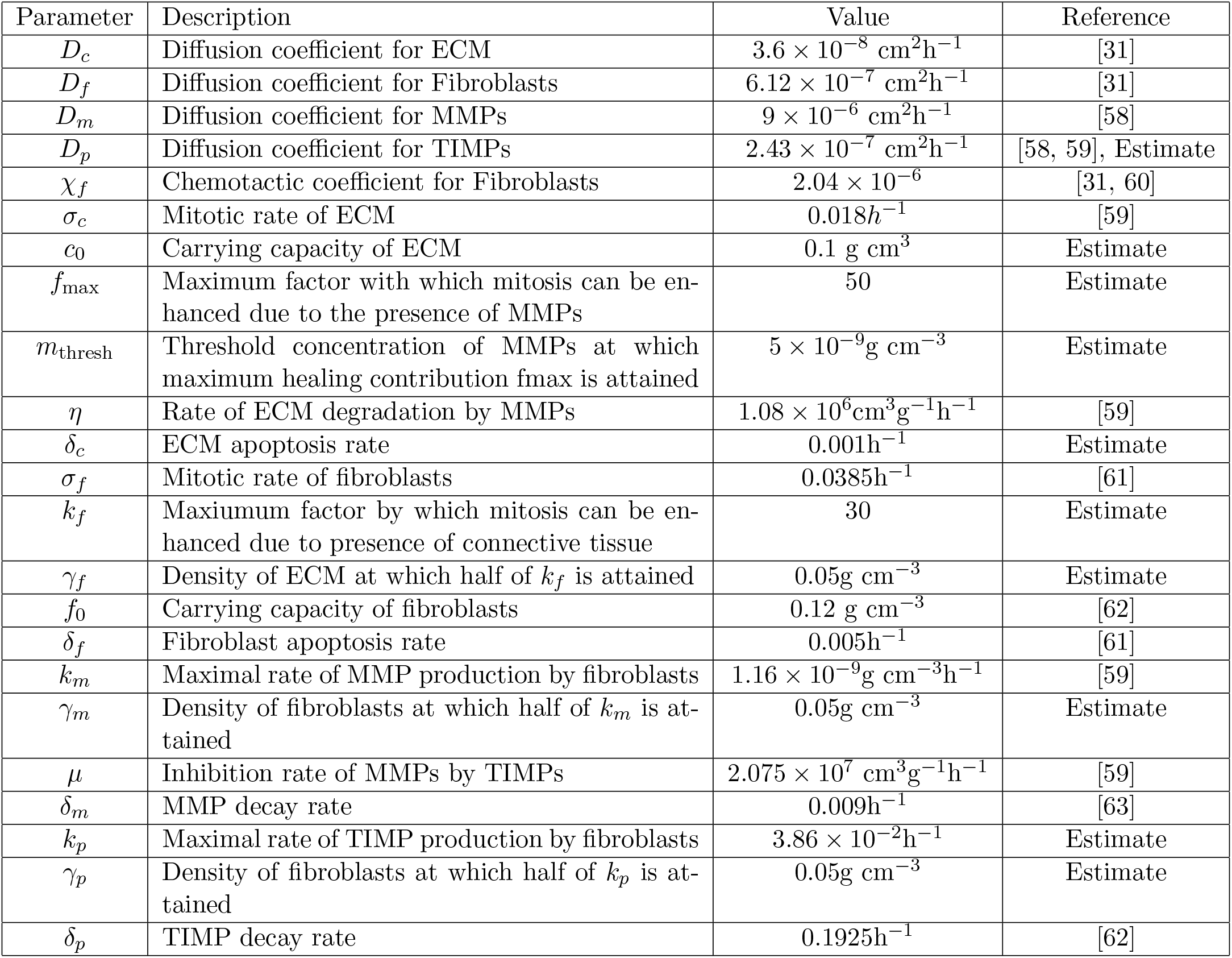
Values and definitions of the dimensional parameters appearing in (1)–(4).

| Parameter | Description | Value | Reference |
| --- | --- | --- | --- |
| $D_c$ | Diffusion coefficient for ECM | $3.6 \times 10^{-8} \text{ cm}^2\text{h}^{-1}$ | [31] |
| $D_f$ | Diffusion coefficient for Fibroblasts | $6.12 \times 10^{-7} \text{ cm}^2\text{h}^{-1}$ | [31] |
| $D_m$ | Diffusion coefficient for MMPs | $9 \times 10^{-6} \text{ cm}^2\text{h}^{-1}$ | [58] |
| $D_p$ | Diffusion coefficient for TIMPs | $2.43 \times 10^{-7} \text{ cm}^2\text{h}^{-1}$ | [58, 59], Estimate |
| $\chi_f$ | Chemotactic coefficient for Fibroblasts | $2.04 \times 10^{-6}$ | [31, 60] |
| $\sigma_c$ | Mitotic rate of ECM | $0.018\text{h}^{-1}$ | [59] |
| $c_0$ | Carrying capacity of ECM | $0.1 \text{ g cm}^3$ | Estimate |
| $f_{\max}$ | Maximum factor with which mitosis can be enhanced due to the presence of MMPs | 50 | Estimate |
| $m_{\text{thresh}}$ | Threshold concentration of MMPs at which maximum healing contribution $f_{\max}$ is attained | $5 \times 10^{-9} \text{ g cm}^{-3}$ | Estimate |
| $\eta$ | Rate of ECM degradation by MMPs | $1.08 \times 10^6 \text{ cm}^3 \text{g}^{-1} \text{h}^{-1}$ | [59] |
| $\delta_c$ | ECM apoptosis rate | $0.001\text{h}^{-1}$ | Estimate |
| $\sigma_f$ | Mitotic rate of fibroblasts | $0.0385\text{h}^{-1}$ | [61] |
| $k_f$ | Maximum factor by which mitosis can be enhanced due to presence of connective tissue | 30 | Estimate |
| $\gamma_f$ | Density of ECM at which half of $k_f$ is attained | $0.05 \text{ g cm}^{-3}$ | Estimate |
| $f_0$ | Carrying capacity of fibroblasts | $0.12 \text{ g cm}^{-3}$ | [62] |
| $\delta_f$ | Fibroblast apoptosis rate | $0.005\text{h}^{-1}$ | [61] |
| $k_m$ | Maximal rate of MMP production by fibroblasts | $1.16 \times 10^{-9} \text{ g cm}^{-3} \text{h}^{-1}$ | [59] |
| $\gamma_m$ | Density of fibroblasts at which half of $k_m$ is attained | $0.05 \text{ g cm}^{-3}$ | Estimate |
| $\mu$ | Inhibition rate of MMPs by TIMPs | $2.075 \times 10^7 \text{ cm}^3 \text{g}^{-1} \text{h}^{-1}$ | [59] |
| $\delta_m$ | MMP decay rate | $0.009\text{h}^{-1}$ | [63] |
| $k_p$ | Maximal rate of TIMP production by fibroblasts | $3.86 \times 10^{-2} \text{ h}^{-1}$ | Estimate |
| $\gamma_p$ | Density of fibroblasts at which half of $k_p$ is attained | $0.05 \text{ g cm}^{-3}$ | Estimate |
| $\delta_p$ | TIMP decay rate | $0.1925\text{h}^{-1}$ | [62] |

**Table 2:** Values and definitions of the dimensionless parameter values appearing in the Regulated Wound Healing Model in (11)–(19).

| Parameter | Definition | Value |
| --- | --- | --- |
| $\sigma_C$ | $\frac{f_0}{c_0}$ | 1.2 |
| $\bar{\eta}$ | $\frac{\eta m_{\text{thresh}}}{\sigma_c}$ | 0.30 |
| $\delta_C$ | $\frac{\delta_c}{\sigma_c}$ | 0.056 |
| $D_F$ | $\frac{D_f}{D_c}$ | 17 |
| $\chi_F$ | $\frac{\chi_f}{D_c}$ | 57 |
| $\sigma_F$ | $\frac{\sigma_f}{\sigma_c}$ | 2.1 |
| $\gamma_F$ | $\frac{\gamma_f}{c_0}$ | 0.50 |
| $\delta_F$ | $\frac{\delta_f}{\sigma_c}$ | 0.28 |
| $D_M$ | $\frac{D_m}{D_c}$ | 250 |
| $k_M$ | $\frac{k_m}{m_{\text{thresh}}\sigma_c}$ | 13 |
| $\gamma_M$ | $\frac{\gamma_m}{f_0}$ | 0.42 |
| $\bar{\mu}$ | $\frac{\mu m_{\text{thresh}}}{\sigma_c}$ | 5.8 |
| $\delta_M$ | $\frac{\delta_m}{\sigma_c}$ | 0.50 |
| $D_P$ | $\frac{D_p}{D_c}$ | 6.8 |
| $k_P$ | $\frac{k_p}{\sigma_c}$ | 2.1 |
| $\gamma_P$ | $\frac{\gamma_p}{f_0}$ | 0.42 |
| $\delta_P$ | $\frac{\delta_p}{\sigma_c}$ | 11 |
| $\bar{L}$ | - | 170 |
| $\bar{\kappa}$ | - | 0.1 |
| $\lambda$ | - | 3 |

### 2.2 Model Simulations

In order to obtain numerical simulations of the RWHM, we discretise the one-dimensional spatial domain [0, *L*] into *N*_*x*_ = 1000 equally-spaced nodes, using second-order finite differences to approximate spatial derivatives. The resulting system of time-dependent ODEs was then integrated using MATLAB’s ODE15s function.

Subject to the parameter values in Table 2, which represent well-regulated wound healing, we obtain the numerical solutions of the RWHM in Figure 2. Here, we observe the evolution of ECM density *C*(*X, T*) into a travelling wave, where a state close to unity, (i.e. a healed state), invades a wound (*C* = 0). The invasion of a wound by a healed front of tissue suggests the complete healing of an acute wound. We also observe the formation of travelling waves for fibroblasts, *F* (*X, T*), where their invading state is close to unity. Additionally, during the wound healing process, we observe travelling waves for MMPs, *M* (*X, T*) and TIMPs, *P* (*X, T*), as shown in Figures 2(c) and (d) respectively. Notably, a spike in the concentrations of MMPs and TIMPs is observed at the edge of the healing wound. We note that this spike occurs naturally and arises across a broad range of initial conditions. These observations are consistent with the biological literature [5, 10], which suggest that there is an increased expression of MMP concentration at the edge of a healing wound in all stages of the healing of an acute wound.

**Figure 2:**
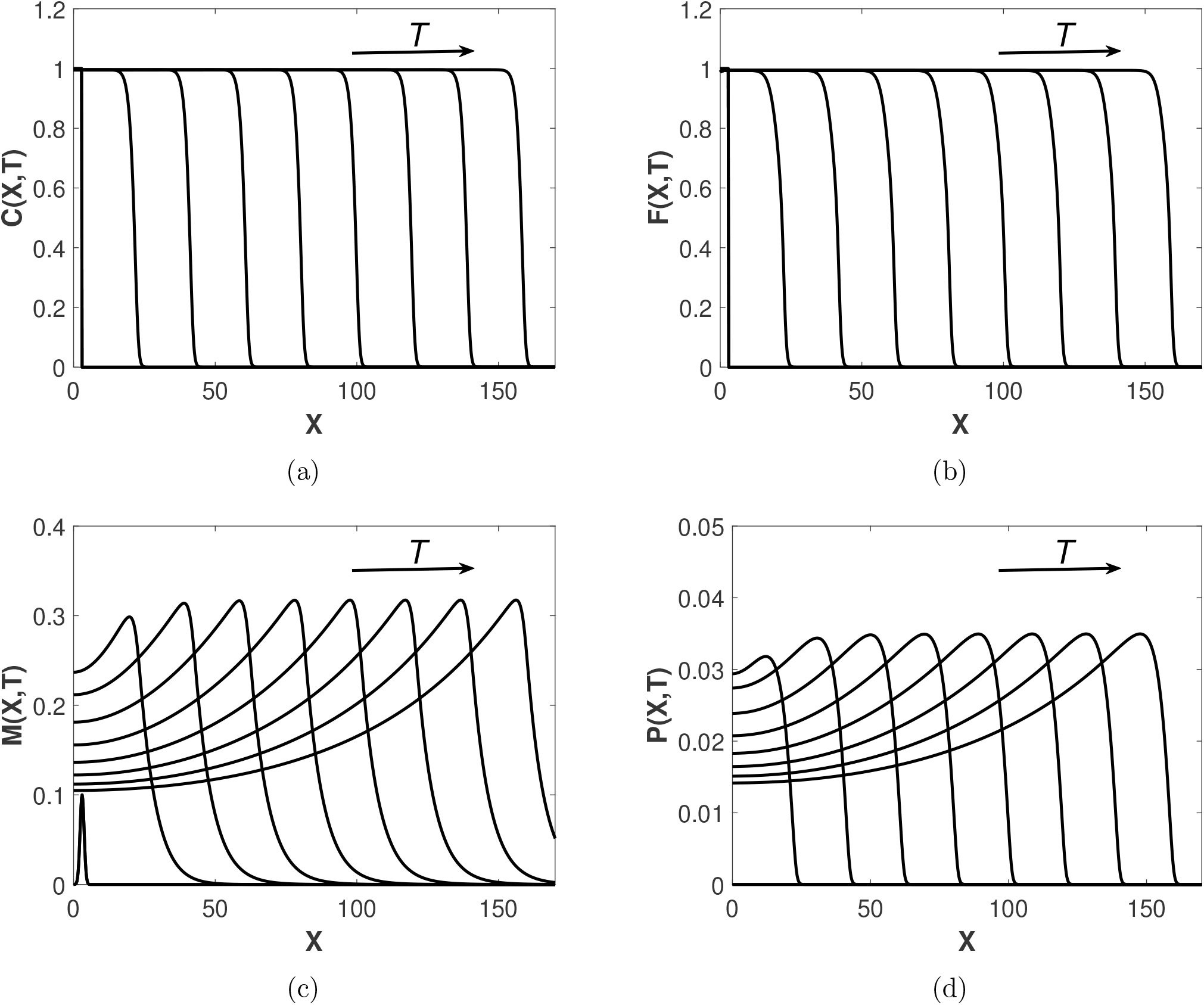
Simulation of the RWHM (11)–(19) at regular time intervals *T* = 0.5 with all parameter values as in Table 2. Evolution of (a) ECM density *C*(*X, T*), (b) fibroblast density *F* (*X, T*), (c) MMP concentration *M* (*X, T*) and (d) TIMP concentration *P* (*X, T*) across the spatial domain.

## 3 Steady States and Bifurcation Analysis

From the numerical solutions of the RWHM seen in Figure 2, we observe the emergence of travelling wave solutions. Therefore, in this section, we examine the uniform steady states of our system corresponding to the far-field states of the travelling wave solutions. In particular, the non-zero far-field ECM density, 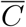, will determine whether or not a wound heals to completion. The steady states of the system 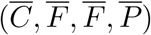 are given by setting the temporal and spatial derivatives in the RWHM to zero:

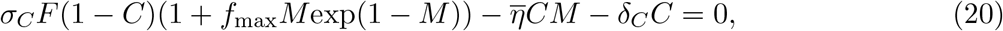

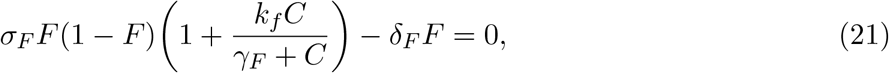

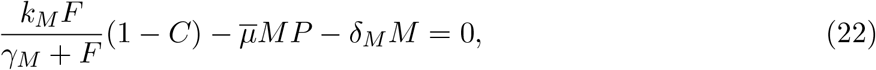

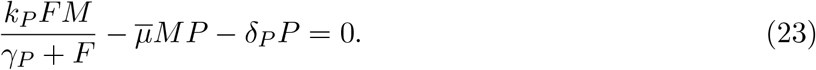

While the above equations cannot be solved analytically for non-trivial solutions, it is clear that we always have the trivial steady state (*C, F, M, P*) = (0, 0, 0, 0) for all parameter sets.

The biological literature suggests that one potential cause for the emergence of chronic wounds is deregulated apoptosis, which may be caused by uncontrolled blood sugars in diabetes patients [13]. Therefore, we numerically solve the steady state equations to determine 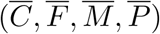 and examine how 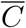 changes with the scaled apoptotic rate of the ECM, *δ*_*C*_. Bifurcation diagrams are presented in Figure 3, where we note the presence of a bistable region bounded by two saddle-node bifurcation points. This phenomenon indicates that if *δ*_*C*_ is increased sufficiently, the invading state of the travelling waves will transition to the lower, unhealed branch. In [18], a similar bifurcation diagram was presented, showing how the steady states of dermal cells change with *δ*_*C*_, which also exhibited the existence of such a bistable region. It was demonstrated that if the invading state takes a value on the lower chronic wound branch, then a successful treatment will require substantial reduction in *δ*_*C*_ so as to pass beneath the bistable region and transition back up to the healed branch, meaning that the wound is more difficult to heal, and demonstrating the presence of a hysteresis loop [64]. This loop indicates the persistence of multi-stable travelling waves, depending on initial conditions corresponding to either a healed or chronic wound [18]. These dynamics were demonstrated through numerical simulations, providing valuable insights into the maintenance of chronic wounds.

**Figure 3:**
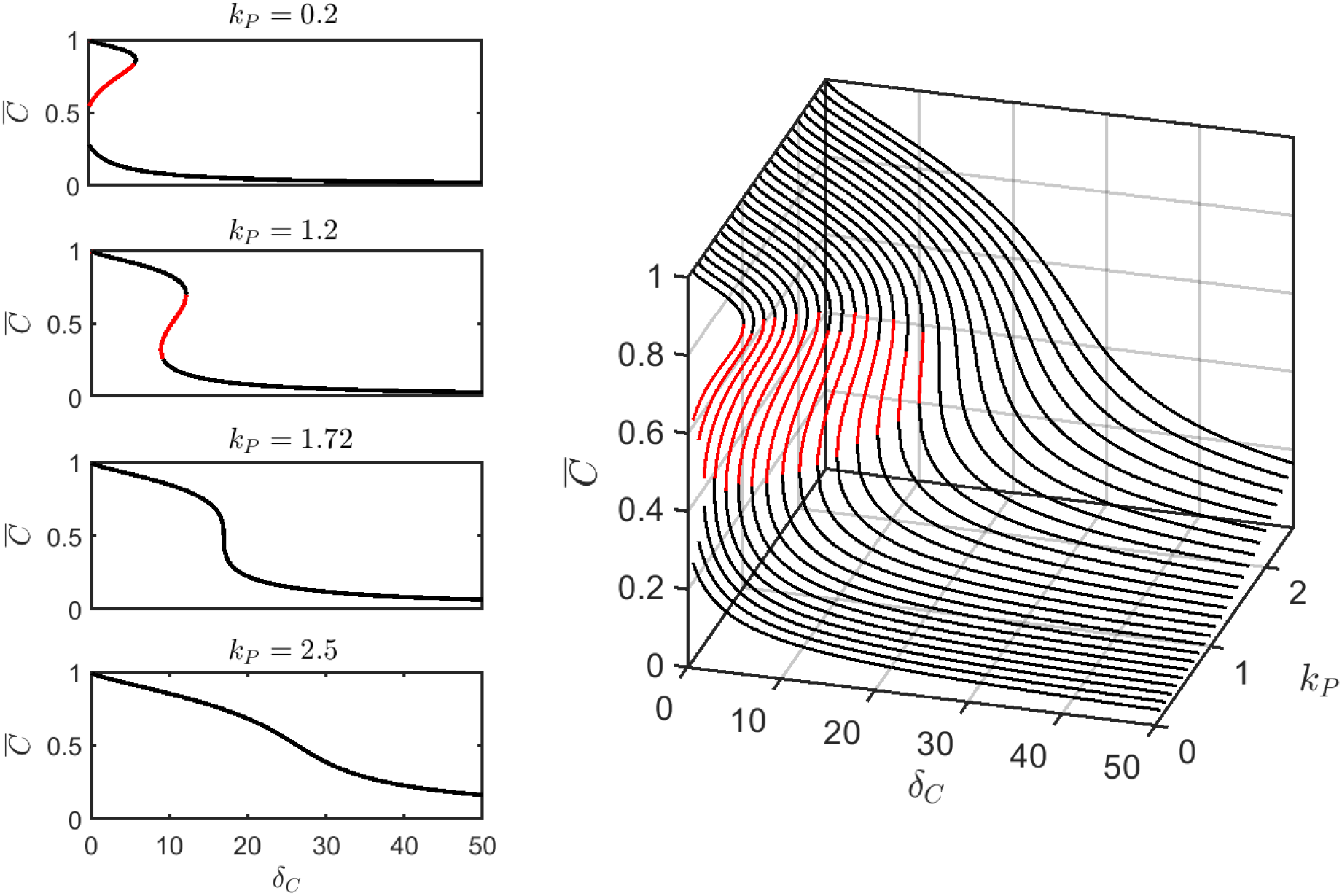
Bifurcation diagrams of (20)–(23) showing the behaviour of the steady state 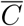 under variation of *δ*_*C*_ and *k*_*P*_. The solid black lines represent stable branches and the solid red lines represent unstable branches. We also display bifurcation diagrams in the 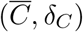-plane for specified values of *k*_*P*_.

We now extend this bifurcation analysis by examining the influence of TIMP production, *k*_*P*_ on the bifurcation diagrams for *δ*_*C*_ in Figure 3. We observe that for small TIMP production, one of the saddle-node bifurcation points exists in the negative *δ*_*C*_ region, therefore, the invading state of the travelling wave cannot return to the healed branch from the lower chronic wound branch in the physically relevant case of positive *δ*_*C*_. As a result, the model predicts that for low TIMP production, chronic wounds cannot be reversed due to insufficient MMP regulation, making external treatment essential for restoring the wound to a healed state. As *k*_*P*_ is increased, we observe dynamics similar to that of the model in [18] – a hysteresis loop in which it is possible to transition from a chronic wound to a healed wound by decreasing *δ*_*C*_ sufficiently enough such that we are able to return to the upper healed branch. As *k*_*P*_ is increased further, the unstable branch is destroyed, resulting in a single stable branch. The regulatory potential of *k*_*P*_ and *δ*_*C*_ on the healing of the ECM is summarised by the two-parameter continuation of *δ*_*C*_ against *k*_*P*_ in Figure 4. From this figure, we deduce that below the value *k*_*P*_ = 0.31, we observe irreversible chronic wounds where one of the saddle-node bifurcation points exists in the negative *δ*_*C*_ region, while for values above *k*_*P*_ = 1.71, the hysteresis loop is destroyed and we see only a single stable non zero branch for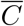. For values of *k*_*P*_ between the above two threshold values, we identify a hysteresis loop representing reversible chronic wounds. This bifurcation analysis suggests that the TIMP production rate significantly contributes to the regulation of the healing of chronic wounds as a result of its MMP inhibiting properties.

**Figure 4:**
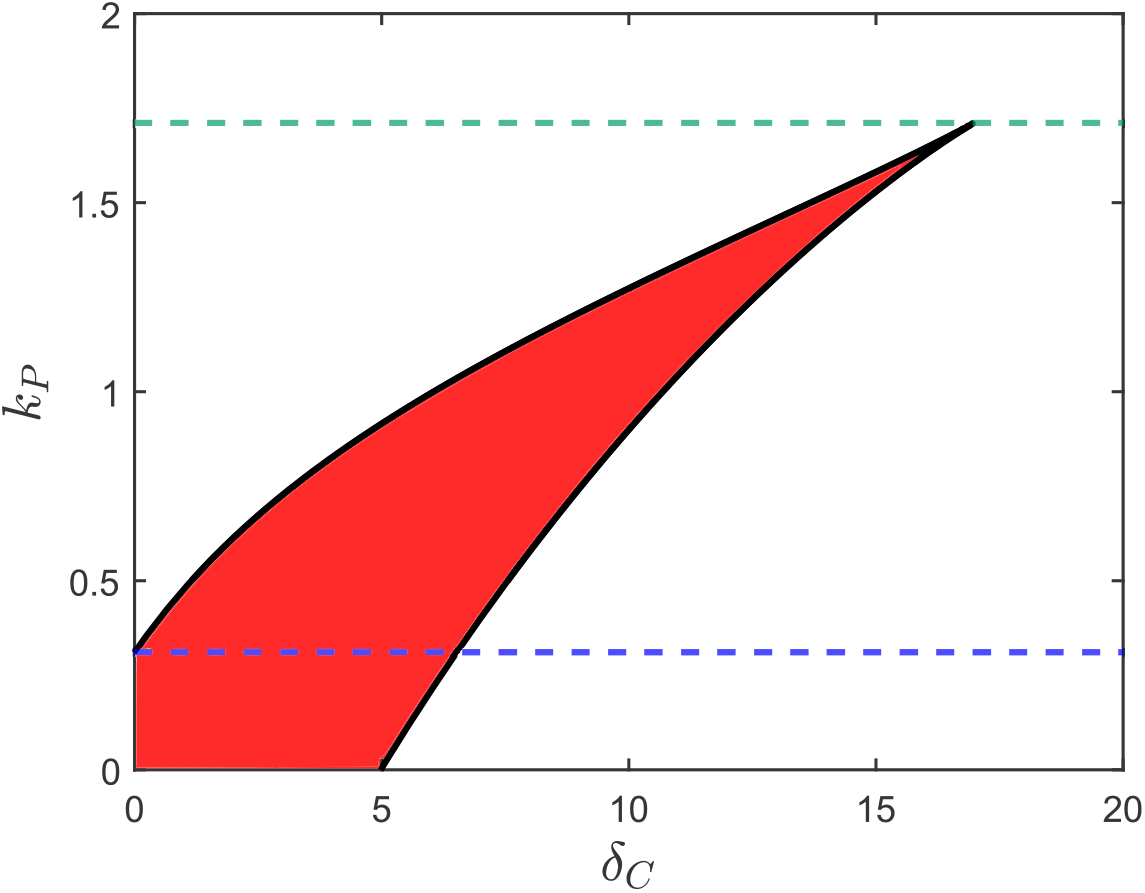
Two-parameter continuation of the interaction of the apoptotic rate of ECM parameter, *δ*_*C*_ and the TIMP production parameter, *k*_*P*_. The red highlighted region represents the parameter space where bistability is observed. The blue and green dashed lines, at *k*_*P*_ = 0.31 and *k*_*P*_ = 1.71 respectively, are the thresholds of changing qualitative behaviours.

## 4 Parameter Sensitivity and Model Reduction

In this section, we conduct a parameter sensitivity analysis of the RWHM. Currently, there are 19 dimensionless parameters; we seek to retain only those representing the mechanisms of greatest significance. This reduction in the number of parameters allows us to simplify the RWHM and enhance our understanding of the key mechanisms driving wound healing which may aid in developing effective therapeutic strategies. Additionally, reducing the model will make it less computationally expensive, which will be beneficial for more computationally intensive numerical simulations. We adopt the Extended Fourier Amplitude Sensitivity Test (eFAST), which is based on variance decomposition techniques (see [65]). The individual influences of the parameters, as well as their combined interactions on designated sensitivity metrics, are quantified using Fourier analysis. We consider three sensitivity metrics: the far-field state of ECM (the value of 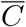 at *x* = 0), the wave speed of the resulting travelling waves, and the maximum MMP concentration in the ‘spike’. We choose the far-field state of ECM as it determines whether or not a wound heals to completion; the wave speed is indicative of whether the wound is healing at a healthy rate; elevated MMP concentrations may lead to chronic wounds by degrading the ECM and therefore the maximum MMP concentration provides insight into the potential development of chronic wounds.

The eFAST method yields both a first-order sensitivity and a total-order sensitivity. The first-order sensitivity index assesses how much a single parameter contributes to the output variance, whereas the total-order index considers not only the parameter itself but also its interactions with all other model parameters. We plot the eFAST sensitivity analysis of the RWHM for each of the sensitivity metrics specified earlier in Figure 5. We remark that this figure also contains the first- and total-order sensitivity of an additional “dummy” parameter that does not affect the model output. It is used as a control to provide a baseline for comparison with the other parameters. Parameters with a total-order sensitivity index less than or equal to that of the dummy parameter are regarded as not having a significant impact on the sensitivity metric being considered.

**Figure 5:**
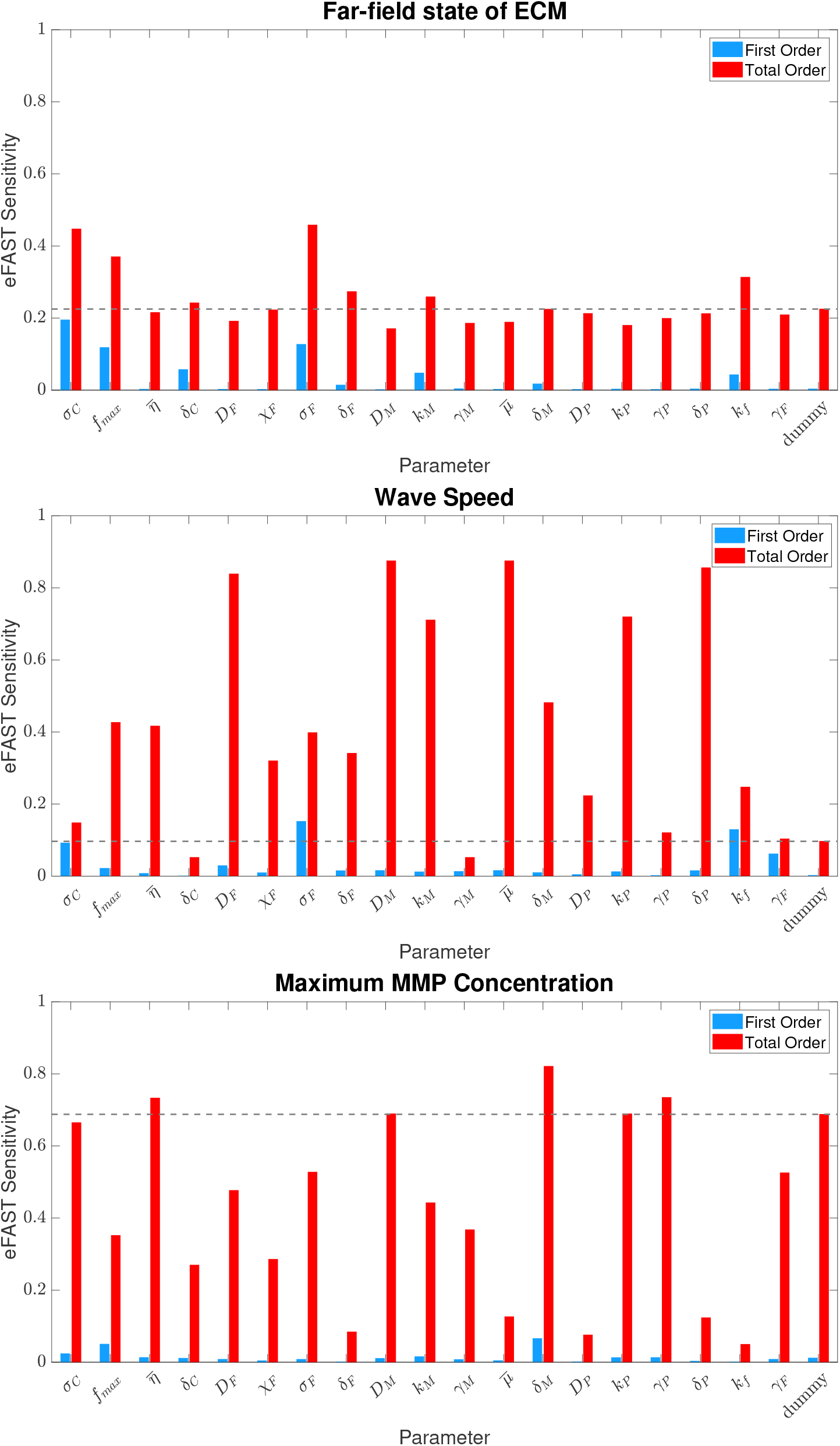
eFAST sensitivity analysis of the parameters of the RWHM on the sensitivity metrics far-field state of the ECM; (b) wave speed and (c) maximum MMP concentration. Blue bars represent the first order sensitivities and red bars represent total order sensitivities. The black dashed line illustrates the total order sensitivity of the dummy parameter. Template code adapted from [66].

From the eFAST sensitivity analysis, we deduce that the parameter groupings that least significantly impact the model outputs of interest are the Michaelis constants, *γ*_*F*_, *γ*_*M*_ and *γ*_*P*_, and the chemotaxis constant *χ*_*F*_. We therefore reduce the model accordingly, by removing the associated processes, to obtain the Regulated Wound Healing Model 2 (RWHM2).

### 4.1 Reduced Model

The reduced RWHM, denoted as the RWHM2, keeps only the most significant parameters in the RWHM and is defined as follows:

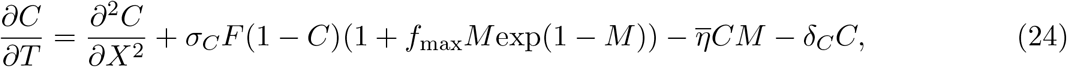

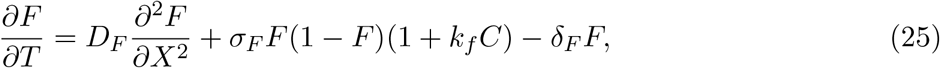

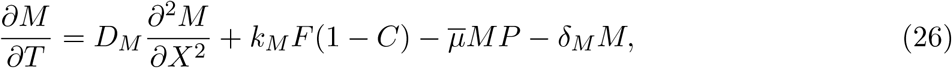

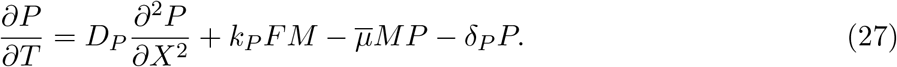

In this subsection, we demonstrate that the RWHM2 displays largely similar qualitative features to the original model.

We present numerical simulations of the RWHM2 in Figure 6 using the same methodology as in Section 2.2 and parameter values as in Table 2. From Figure 6, we observe travelling wave solutions representing the complete healing of an acute wound, i.e. the invasion of a wound by a healed front of cells and ECM. Furthermore, we notice that MMPs and TIMPs also enter the wound with an increased expression of their respective concentrations at the edge of the healing wound. These qualitative observations are consistent with those of the original model as well as the biological literature, as discussed in Section 2.2. In the same way as in Section 3, we plot a two-parameter continuation of the RWHM2; the key qualitative features are consistent with original model shown in Figure 4.

**Figure 6:**
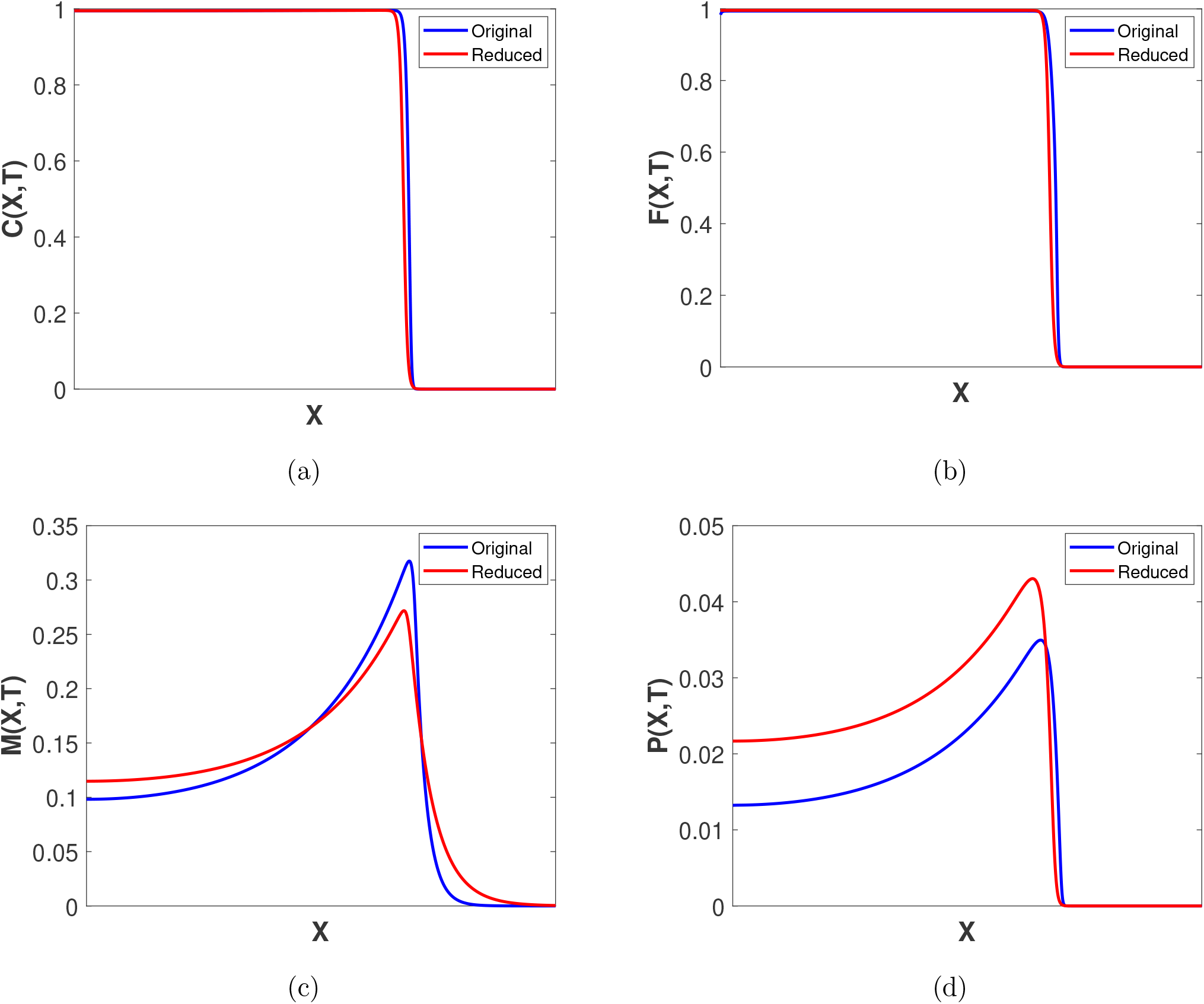
Travelling wave profiles obtained by numerically simulating the original RWHM (11)–(19) in blue, and the reduced RWHM2 in red each at a different time snapshot. The specific time points selected are not critical, as the wave shape remains constant once the traveling wave stabilizes. The comparison is presented for illustrative purposes.

**Figure 7:**
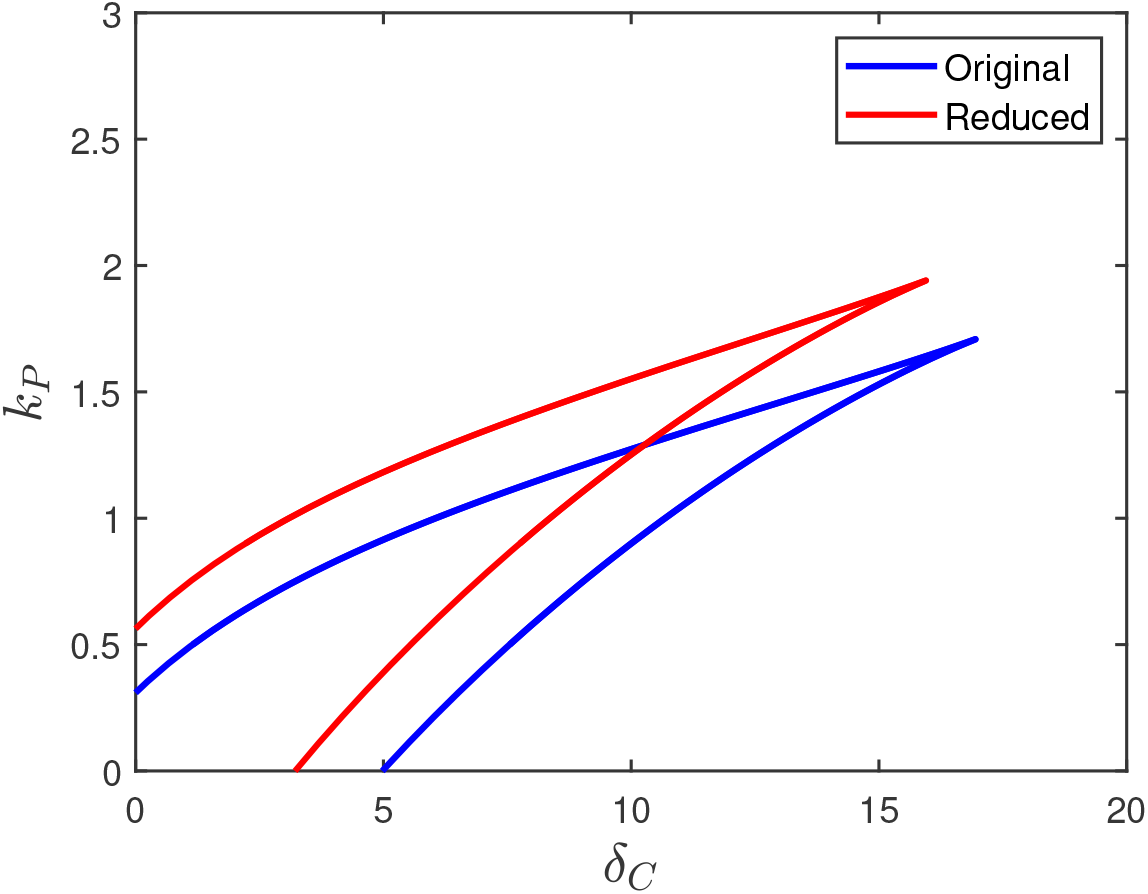
Two-parameter continuation showing the interaction of the apoptotic rate of ECM parameter, *δ*_*C*_ and the TIMP production parameter, *k*_*P*_ for the RWHM in blue, and the RWHM2 in red. All other parameters are in Table 2.

From the above analysis, we deduce that the reduced model is able to successfully emulate key qualitative features of the original model while being less computationally expensive (a runtime of 2.98 seconds compared with 7.12 seconds for the original RWHM). We remark that the precise shapes of the waves and the locations of behavioural shifts in parameter space are somewhat arbitrary, as we have not performed any fitting to experimental data. Therefore, these minor deviations between the two models are not significant. This reduction in computational cost is particularly advantageous when extending the model to a two-dimensional spatial domain as we will do in Section 5, since it allows us to simulate wound healing more efficiently without sacrificing biological realism.

## 5 Two-Dimensional Model Simulations

In Section 3, we determined that increased TIMP production results in better regulation of chronic wounds. However, altering the intrinsic TIMP production is not a practical treatment approach. Instead, a more feasible treatment may involve applying an external source of TIMPs to the wound by means of a hydrogel – an approach currently at the research phase [67, 68]. Therefore, in this section, we extend the RWHM2 into two dimensions to simulate a more biologically realistic wound, allowing us to study the treatment of an external TIMP source on the wound surface. First we consider the effect of elevated MMP production on the healing of a wound with no treatment via TIMPs which leads to the emergence of a chronic wound. We then consider this chronic wound and model the application of an external source of TIMPs to the wound.

The dimensionless RWHM2 posed on a two-dimensional spatial domain 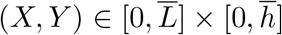 is as follows:

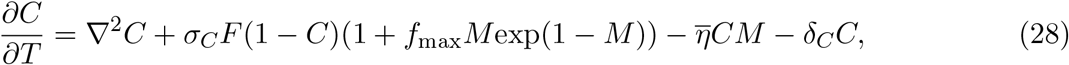

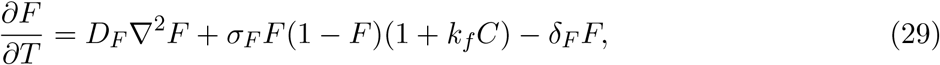

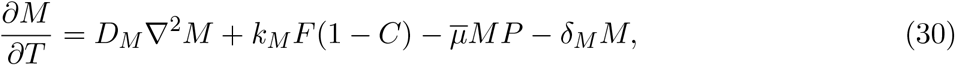

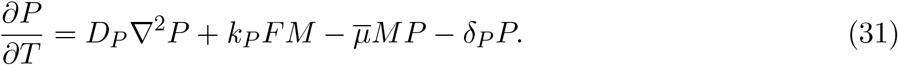

The non-dimensionalisation is obtained using the same scalings in Section 2.1, with the *y*-coordinate scaled as 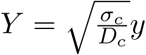. We note that the 2D equations (28)–(31) with analogous initial and boundary conditions and parameter values to those in Section 2.1 yield travelling waves with no *y*-dependence. These waves exhibit the same qualitative features and wave speed as the 1D reduced model, thereby affirming the validity of the 2D numerical approximations that we employ (see Appendix A).

p We now examine the impact of an elevated value of *k*_*M*_, the MMP production parameter, on wound healing in our 2D model, in the presence and absence of treatment by an external source of TIMPs. In the first case of no treatment, we adopt zero flux boundary conditions as in Section 2.2; treatment via an external source of TIMPs applied to the wound surface is modelled via a Robin boundary condition on the top edge of the 2D wound:

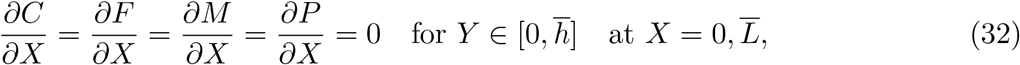

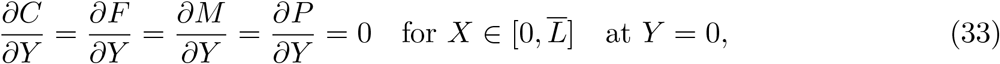

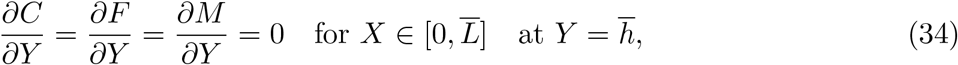

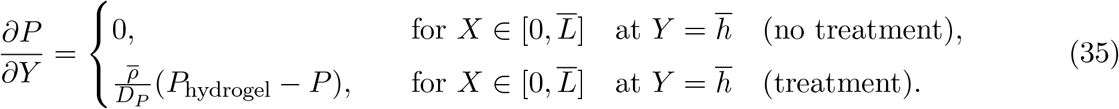

The procedure for deriving this non-dimensionalised Robin boundary condition is detailed in Appendix B. To assess wound closure as an indicator of healing, we consider a wound that is initially symmetric about the centreline 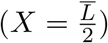, resulting in the following initial conditions:

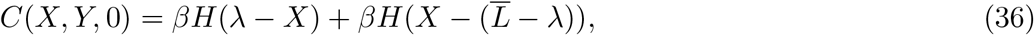

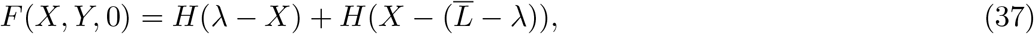

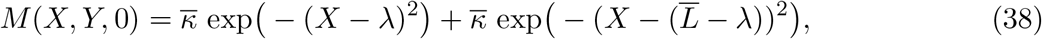

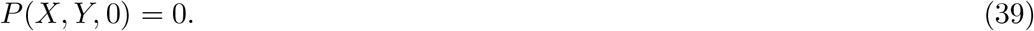

We note that the chosen values of *β* and 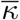 are different in the two cases that we will consider. In the case of no treatment, we use *β* = 1 and 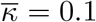 as in the results presented in Section 2 in the case of the 1D RWHM. Later, we demonstrate that no treatment and elevated MMP production leads to travelling waves representing the emergence of a chronic wound. Therefore, in the case of treatment via an external source of TIMPs, we start with the chronic wound from the untreated case, using *β* = 0.05 and 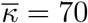 to ensure the development of travelling waves characteristic of the chronic wound by matching the far-field state for the ECM and the peak MMP concentration.

In order to obtain numerical simulations of the 2D RWHM2, we use MATLAB’s PDE toolbox and consider a slender domain where 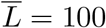 and 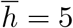. This aspect ratio allows the 2D numerical simulations to be well approximated by the 1D RWHM2 as expected, provided that analogous zero-flux boundary conditions (but not Robin boundary conditions) are applied. Firstly, we present numerical solutions of the 2D RWHM2 (28)–(39) in Figure 8 in the case of elevated MMP production levels and no treatment. Figure 8 illustrates the development of travelling waves representing chronic wounds that are independent of *Y*. In these simulations, the invading state for *C*(*X, Y, T*) maintains an unhealed value, while the wound is characterised by elevated MMP concentrations. Eventually, *C*(*X, Y, T*) settles at this unhealed value and *M* (*X, Y, T*) settles at a large value which we characterise as an unhealthy, chronic wound state. We now aim to investigate the therapeutic potential of a hydrogel containing TIMPs administered to a chronic wound, characterised by a Robin boundary condition on the top edge of the 2D wound as given in (35). We present illustrative simulations of the 2D RWHM2 in Figure 9 to show the treatment of a chronic wound with an external source of TIMPs. The travelling waves for *C*(*X, Y, T*) settle at a healed state 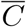, as the additional TIMPs effectively regulate the elevated MMP concentrations, causing *M* (*X, Y, T*) to settle at a reduced and healthy value 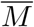. This regulation ultimately allows the wound to heal fully. We also observe that all variables, with the exception of *P* (*X, Y, T*), exhibit minimal variation in the *Y* -direction and therefore we display a comparison of the *Y* -averaged travelling waves for *C*(*X, Y, T*) presented in Figures 8 and 9 in Appendix C. These results align with the biological literature, indicating that TIMPs regulate elevated MMP concentrations and promotes better healing of chronic wounds. Our results therefore support the use of hydrogels containing TIMPs as an effective treatment for chronic wounds. While this study provides a proof of concept, future research should focus more closely on the specific physical and chemical components of such hydrogels to better inform their development and application in treatments.

**Figure 8:**
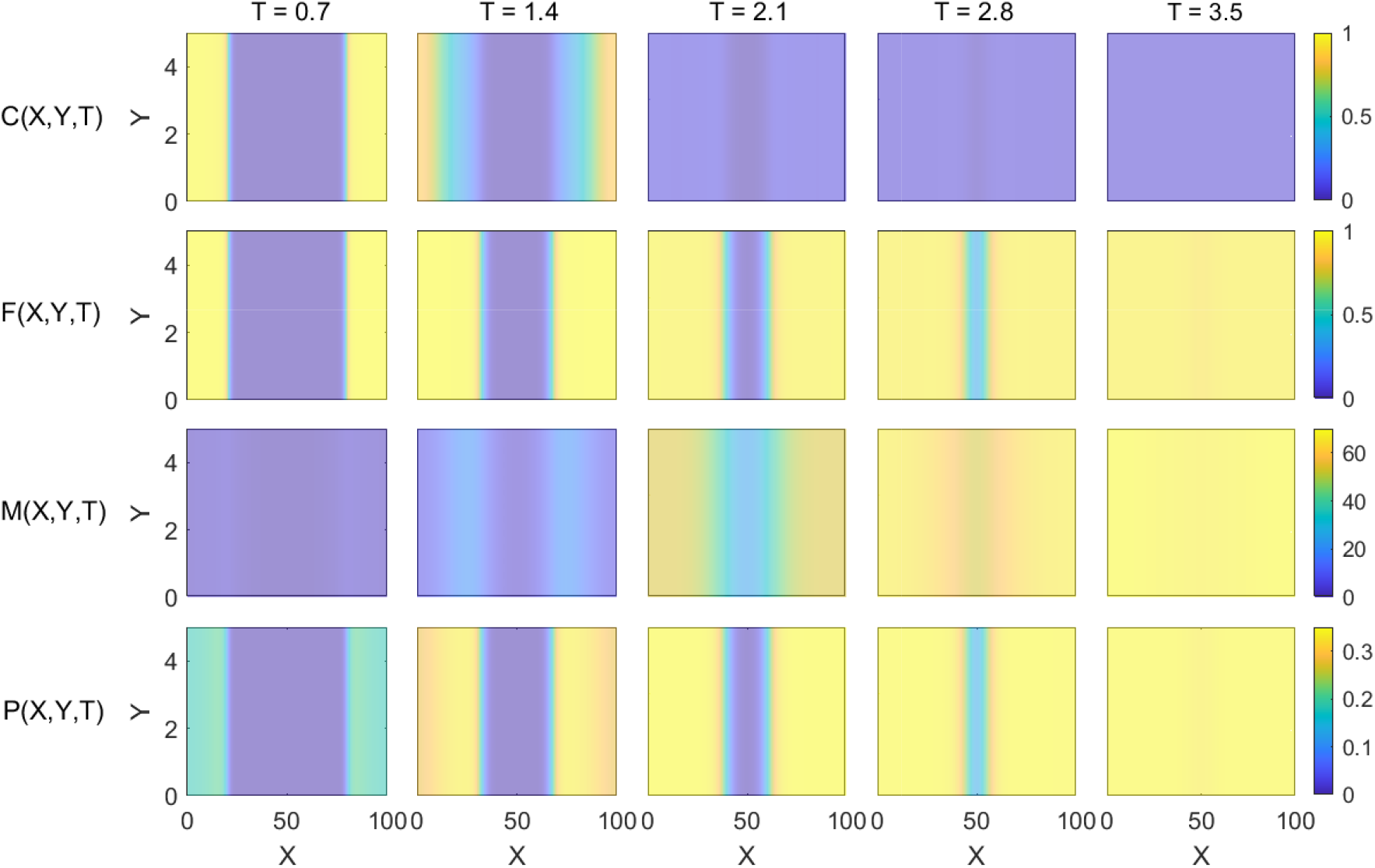
Numerical solutions of the 2D RWHM2 (28)–(39) at regular intervals *T* = 0.7. 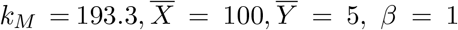, and 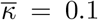. All other parameter values as are as in Table 2. Initial conditions as in (36)–(39) representing a symmetric synthetic wound and zero flux boundary conditions as in (32)–(35)

**Figure 9:**
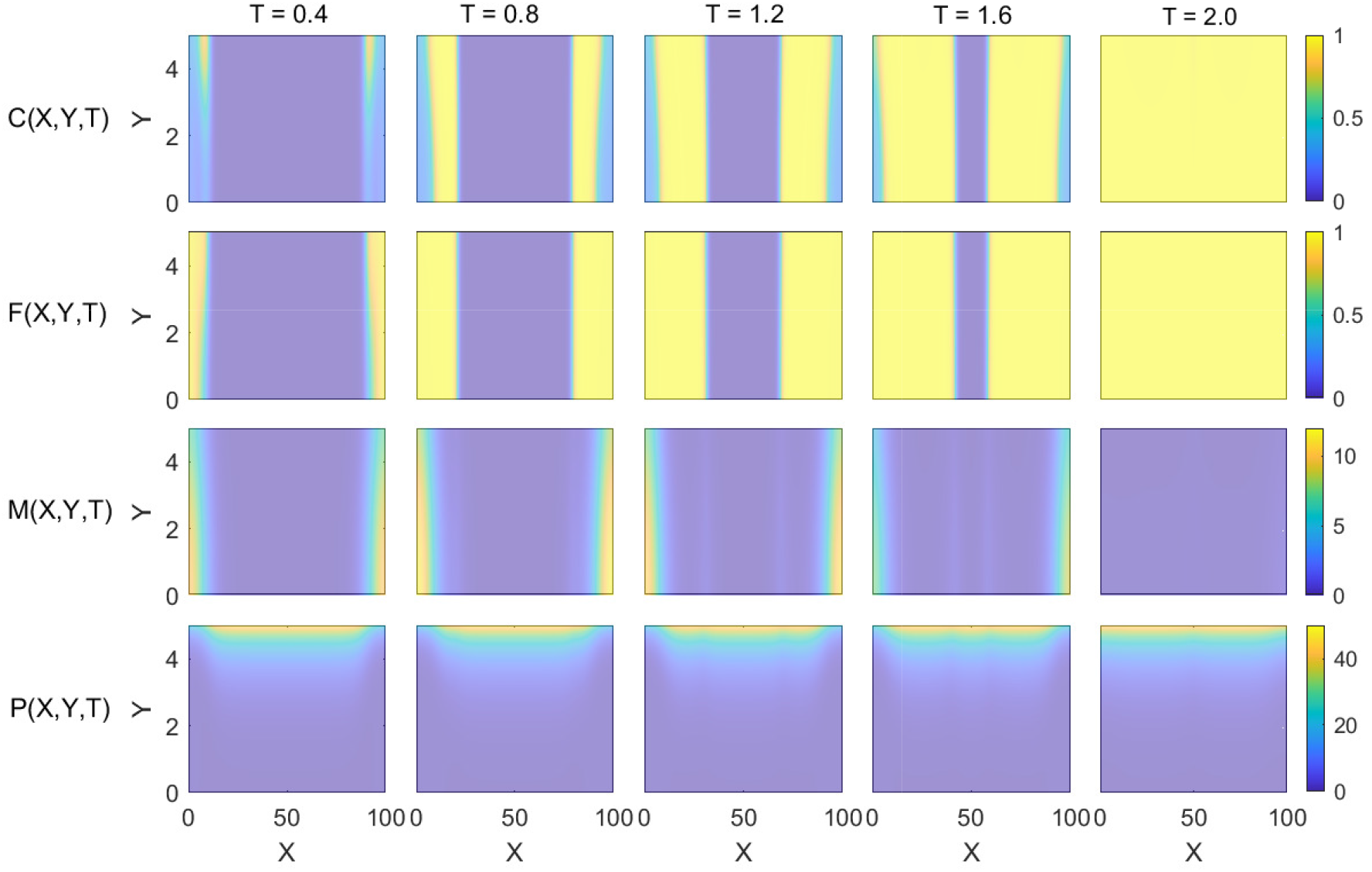
Numerical solutions of the 2D RWHM2 (28)–(39) at regular intervals *T* = 0.4. 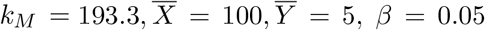 and 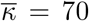. All other parameter values as are as in Table 2. Initial conditions as in (36)–(39) representing a symmetric synthetic wound and zero flux boundary conditions as in (32)–(35), noting that we apply the Robin boundary condition as defined in (35), with 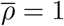 and *P*_hydrogel_ = 445. We also note the colourbar limits change for *P* (*X, Y, T*) compared with that in Figure 8.

## 6 Conclusions and Discussion

MMPs and TIMPs both play a crucial role in the development and management of chronic wounds. Elevated MMP concentrations contribute to the development of chronic wounds as they destroy the surrounding ECM, while TIMPs regulate MMP activity, thereby regulating wound progression. Therefore, in this work, we develop a system of reaction-diffusion equations to describe the interaction of MMPs and TIMPs in both acute and chronic wounds. Our mathematical model gives rise to travelling wave solutions and is able to replicate key qualitative behaviours observed physiologically. One such property is the increased expression of MMP concentrations at the edge of a healing wound across all stages of acute wound recovery. Furthermore, biological literature suggests that one factor contributing to chronic wound formation is deregulated apoptosis. Therefore, we explore the impact of varying the apoptotic rate of the ECM across different levels of TIMP production to investigate the regulatory effects of TIMPs. Using bifurcation analysis, we observe a region of bistability which indicates that that small TIMP production levels lead to irreversible chronic wounds, while increased TIMP production levels result in reversible chronic wounds, which are better regulated for larger TIMP production values. These results indicate that high TIMP concentrations lead to more effectively regulated chronic wounds, consistent with existing biological research.

The biologically significant features of this model are retained in a simplified model obtained using an eFAST parameter sensitivity analysis, while reducing computational runtime. This reduced model is further extended to two dimensions to explore the effectiveness of a hydrogel treatment. We find that elevated MMP production levels lead to the emergence of chronic wounds, consistent with biological studies. We consider such chronic wounds and employ a Robin boundary condition to model the application of a hydrogel on the wound surface, which provides an external source of TIMPs. Simulation results demonstrate the reversal of the chronic wound, thereby reaffirming the regulatory properties of TIMPs and supporting the use of TIMP-containing hydrogels as a viable treatment option for chronic wounds. We remark that this method of modelling a hydrogel serves as a proof of concept and future research should focus more on the physical and chemical properties of the hydrogel to develop a more biologically realistic model, which could subsequently be validated by experimental data and thus inform treatment strategies. Additionally, extending the model to more realistic wound geometries may further add to its biological realism.

We note that our study only defines a chronic wound by deregulated apoptosis and elevated MMP production starting from a parameter set corresponding to the healing of an acute wound. However, parameter values for chronic wounds may encompass many variations beyond those considered here which should be considered in future work. Additionally, the presence of a hysteresis loop in the bifurcation diagrams suggests the existence of multistable travelling waves; thus, future work should, therefore, include a stability analysis of these travelling waves to gain deeper insights into their behaviour and implications.

## Acknowledgements

This research did not receive any specific grant from funding agencies in the public, commercial, or not-for-profit sectors.

## Appendix A: 2D numerical simulations and comparison to 1D model

We consider the 2D RWHM2 model (28)–(31) with initial and boundary conditions:

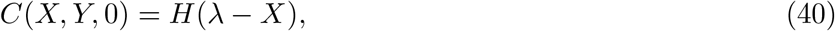

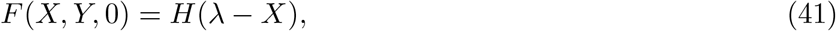

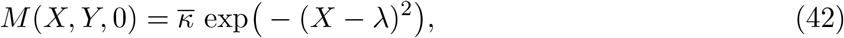

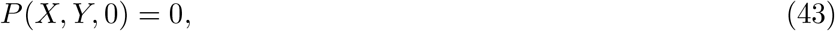

and

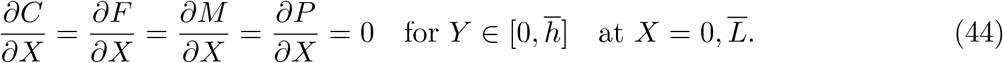

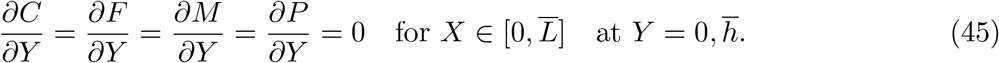

Using the MATLABs PDE toolbox and parameter values in Table 2, we observe travelling wave solutions as shown in Figure 10, representing the complete healing of a wound, i.e. the invasion of a wound by a healed front of cells, with an increased expression of MMP and TIMP concentrations at the edge of the healing wound as in the 1D case. We note that the travelling waves exhibit no *Y* -dependence as expected from the initial and boundary data (40)–(45). Since we only observe variations in the *X*-direction, we also provide the *Y* -averaged travelling waves in Figure 10, generated using MATLAB’s `interpolatesolution` function. This interpolation yields the same wave speed of 24.7 as observed in the 1D case, thereby affirming the validity of the 2D solution.

**Figure 10:**
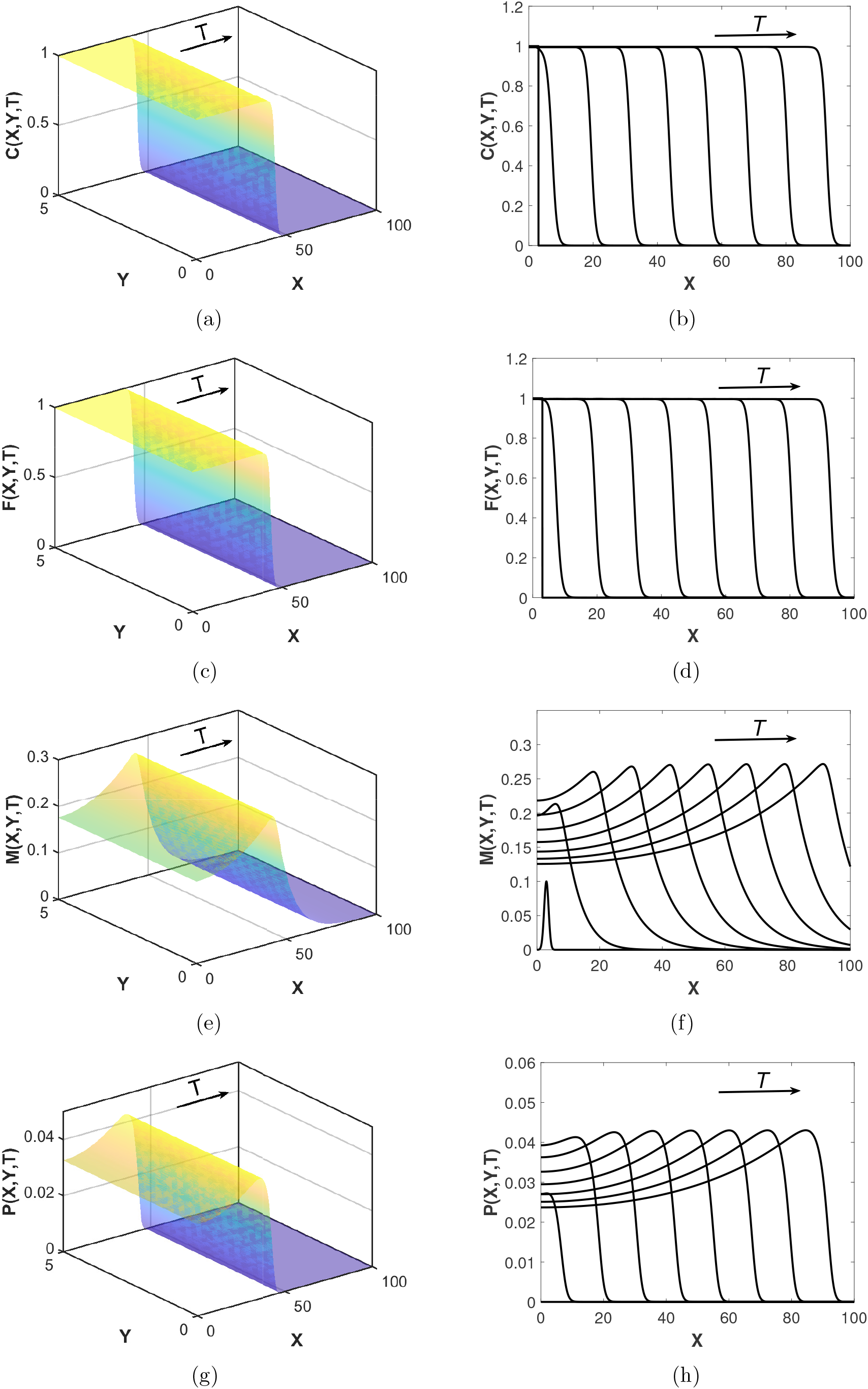
(a) Numerical simulations of the RWHM2 with parameter values as in Table 2 (b) *Y* -averaged travelling waves at time intervals *T* = 0.5 across the *X* domain with parameter values as in Table 2.

## Appendix B: Non-dimensionalisation of the Robin Boundary Condition

The dimensional Robin boundary condition takes the form:

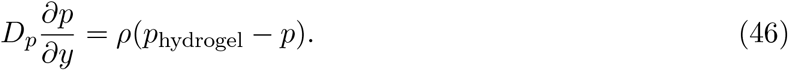

Using the scalings 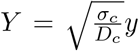 and 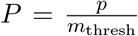, we obtain the non-dimensional Robin boundary condition:

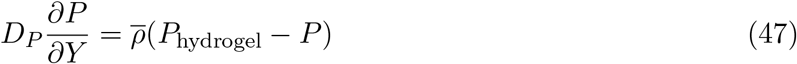

where 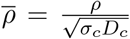 and 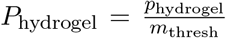 with *ρ* = 2.5456 *×* 10^*−*5^ and *p*_hydrogel_ = 2.225 *×* 10^*−*6^, thereby obtaining 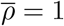.

## Appendix C: *Y* -averaged travelling waves

**Figure 11:**
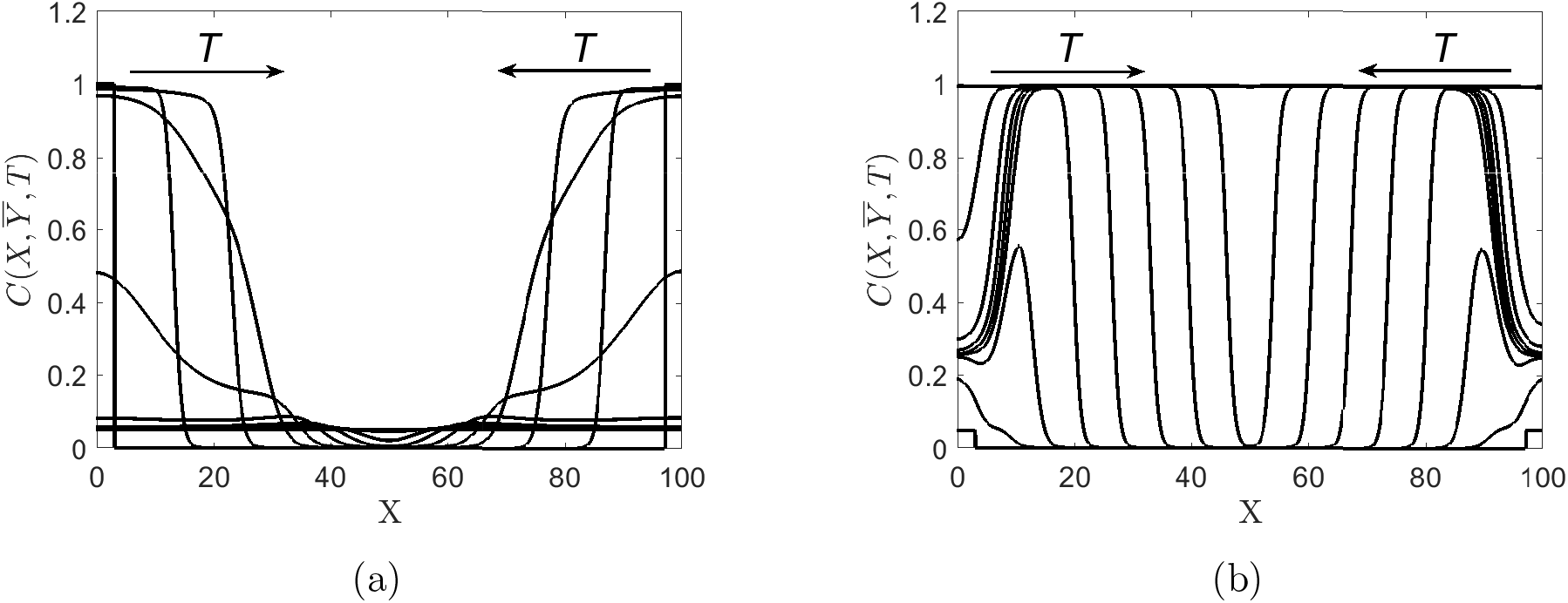
*Y* -averaged travelling waves across the *X* domain with 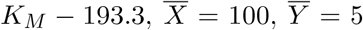 and all other parameter values as in Table 2. (a) Initial conditions as in (36)–(39), with *β* = 1 and 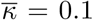 and zero flux boundary conditions as in (32)–(35). (b) Initial conditions as in (36)–(39) with *β* = 0.05 and *κ* = 70 and zero flux boundary conditions as in (32)–(35), except for *P* (*X, Y, T*) on the top edge of the wound, noting that we apply the Robin boundary condition in (35), with 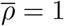 and *P*_hydrogel_ = 445.

